# Termites have developed wider thermal limits to cope with environmental conditions in savannas

**DOI:** 10.1101/2021.05.11.443584

**Authors:** Joel S. Woon, David Atkinson, Stephen Adu-Bredu, Paul Eggleton, Catherine L. Parr

## Abstract

The most diverse and abundant family of termites, the Termitidae, evolved in warm, wet African tropical forests. Since then, they have colonised grassy biomes such as savannas. These environments have more extreme temperatures than tropical forests, and greater temporal fluctuations (both annually and diurnally) that are challenging for soft-bodied ectotherms. We propose that that a likely mechanism that facilitated the expansion from forest to savanna was the widening of physiological limits of savanna termite species in order to cope with more extreme environmental conditions. We sampled termites directly from mound structures across an environmental gradient in Ghana, and recorded the thermal tolerance of individual termites, both critical thermal maximum (CT_max_) and critical thermal minimum (CT_min_). We estimated colony thermal tolerance by taking an average of each tested individual, and modelled these data against several environmental factors (canopy cover above the mound, rainfall and temperature). We found that savanna termite species had significantly higher CT_max_ values, and significantly lower CT_min_ values, than forest species. In addition, areas with high canopy cover were significantly associated with low CT_max_ values, and areas with higher average daily rainfall were significantly associated with higher CT_min_ values. Our results suggest that the widening of thermal tolerances has occurred in savanna termite species, probably in response to the more extreme temperatures found in those environments.

## Introduction

In the terrestrial tropics, there are two dominant biomes: tropical forests and grassy biomes (savannas and grasslands; Mokany, Raison and Prokushkin, 2006; Parr *et al.*, 2014). The spatial distribution of these tropical environments has changed over time, as climatic conditions have varied. Notably, in the late Miocene and into the Pliocene, there was a major expansion of savanna grasslands (Cerling et al., 1993; Cerling et al., 1997; Davies et al., 2020; Pagani et al., 1999), which coincided with the adaptive radiation of C_4_ grasses (Christin et al., 2013; Bouchenak-Khelladi et al., 2014), and a recession of tropical forests. The expansion of savannas offered novel habitats for forest species to colonise, and while this provided opportunity for radiation of species, it also posed specific challenges. Understanding how species radiating from forest to savannas have overcome these challenges is vital for accurately predicting biogeographical history, making inferences of their current ecology, and predicting how these species may be affected by climate change Tropical forests and grassy biomes have markedly different environmental conditions. Forests have greater levels of canopy connectivity, rainfall, humidity (both relative humidity and vapour pressure deficit) and, typically, lower mean temperatures with a smaller thermal range (Nicholls and Wong, 1990; Sternberg, 2001; Tang et al., 2019). In contrast, tropical grassy biomes tend to have more extreme conditions than tropical forests, both spatially with more variable light and therefore temperature regimes, and also temporally over seasonal and diurnal timescales. Specifically, savanna conditions vary annually with pronounced dry seasons and large variation in temperature, humidity and rainfall across the year (D’Onofrio et al., 2019), but also diurnally, with the more open environment allowing for more extreme differences in environmental conditions throughout a 24-hour period (due to higher levels of heat, and insolation, at ground level during the day, and a greater amount of heat loss during the night, when compared with forest; Hardwick et al., 2015).

Despite these environmental differences, a number of taxa have radiated from forests to savannas, adapting to the novel environmental challenges that savannas pose. For example, adaptations of grasses to open tropical environments (Bouchenak-Khelladi et al., 2014; Christin et al., 2013; Pagani et al., 1999) include the notable dominance of C4 grasses in savannas, compared with the prevalence of C3 grasses in tropical forests (Adjorlolo et al., 2012; Still et al., 2014; Solofondranohatra et al., 2018). The generally more open canopy of savannas increases light availability for grass species which, coupled with the increased temperatures and decreased water availability, makes C_4_ photosynthesis more efficient than C_3_ (Still et al., 2014; Solofondranohatra et al., 2018). In contrast, in forests the low understory light levels and higher water availability favour C_3_ photosynthesis over C_4_ (Pearcy and Ehleringer, 1984; Solofondranohatra et al., 2018). Savanna grasses are also commonly taller and more tussocky, which allows them to compete for light more efficiently (Solofondranohatra et al., 2018). In addition, savanna species tend to be harder to decompose, with a higher a C:N ratio, which also facilitates burning (Simon and Pennington, 2012; Solofondranohatra et al., 2018). Forest grass species, on the other hand, tend to be creeping, low-stature species, with large leaves to maximise light intake in the understory, and have structures to minimise self-shading (Solofondranohatra et al., 2018). Closely related plant species can therefore have very different evolutionary strategies to maximise their fitness in different tropical biomes.

Closely related animal species can also have radically different adaptations in different tropical environments. *Bicyclus* butterflies, important pollinators in both tropical African forest and savanna ecosystems (Dierks and Fischer, 2008), evolved a suite of traits when they colonised savannas from tropical forests, including diapause during the dry season, probably triggered by the lack of available suitable surfaces (grasses) for oviposition, which is in contrast to the forest species of *Bicyclus*, which do not have a period of diapause and are able to reproduce all year round (Halali et al., 2020).

However, little is known about how other invertebrate groups may have adapted to the novel environment that savannas present. The Termitidae, the most abundant and diverse group of termites (henceforth all mention of termites refers to the Termitidae family), also evolved inside tropical forests in central Africa (Eggleton, 2000; Aanen and Eggleton, 2005; Bourguignon et al., 2015). Since then they have radiated from forests into more open tropical environments (Aanen and Eggleton, 2005). Forests typically have higher generic and species richness than savannas, although some genera occur in both habitats. However, despite these shared genera, there are almost no shared species between the two environments (Evouna Ondo et al., unpublished data), suggesting that while genera have radiated into savannas, they have since speciated and adapted to savanna environments. Similar to *Bicyclus* butterflies, savanna environments would have posed new challenges to colonising termite species. Termites are soft-bodied ectotherms, and extremely susceptible to desiccation (Heath et al., 1971; Willmer, 1982), and would be challenged by the higher temperatures and lower humidity in savannas. Yet, despite the more extreme conditions, termites have colonised, diversified, and become ecologically dominant in savannas, with a comparable level of biomass to large ungulates or elephants (Dangerfield et al., 1998; Støen et al., 2013). Termites are the dominant soil invertebrate across tropical ecosystems, contributing to multiple ecosystem processes including decomposition, nutrient cycling and bioturbation (Bottinelli et al., 2015; Fox-Dobbs et al., 2010; Joseph et al., 2014; Jouquet et al., 2016; Turner, 2019). As ecosystem engineers, they influence these processes at landscape scales (Davies et al., 2014; Davies et al., 2016a), altering the spatial distribution of nutrients and, as a result, plant species distributions (Davies et al., 2016b; Davies et al., 2016a; Joseph et al., 2014; Støen et al., 2013).

How termites have been able to colonise and to adapt to savanna conditions is currently uncertain. Termites have very different life history traits compared with other invertebrates (Poissonnier et al., 2018; Porter and Hawkins, 2001; Roisin, 2000); most termite species produce complex nest structures, and/ or live underground (Jones and Oldroyd, 2006; Noirot and Darlington, 2000). These subterranean environments could protect and buffer termites from extreme environmental conditions, maintaining stable and ideal nest conditions (Gouttefarde et al., 2017; Jones and Oldroyd, 2006; Korb, 2003). The adaptation of nest structures to provide more efficient thermoregulatory properties, such as altering the external shape to interact with the external temperature to maintain more stable internal conditions (Ocko et al., 2017; Singh et al., 2019; Vesala et al., 2019), may have helped enable forest termite species to colonise and prosper in savanna environments. This thermoregulation of nests could be particularly important for fungus-growing termites, the subfamily Macroterminae within the termitids, which farm fungus from the genus *Termitomyces* (Wood and Thomas, 1989). Maintaining ideal growing conditions for their fungus symbionts would increase colony fitness, so the adaptation of nest thermoregulatory properties would be vital for the success of Macrotermitinae species (Korb and Linsenmair, 2000; Korb, 2003).

While nest thermoregulation is likely to be important in facilitating the movement from forest to savanna, termites forage outside the controlled environment of their nest, often on the surface, risking exposure to extreme environmental conditions (Legendre et al., 2008; Traniello and Leuthold, 2000). One additional key mechanism facilitating the expansion of termites into savanna is the evolution of wider physiological tolerances. Interspecific variation in thermal limits has been shown to occur across different latitudes in other invertebrate lineages (Diptera, *Drosophila* - Rajpurohit, Parkash and Ramniwas, 2008; Kellermann *et al.*, 2012; various, but dominated by Coleoptera, Hymenoptera and Odonata - Sunday *et al.*, 2014) and with exposure to higher temperatures (Diptera, *Drosophila* - Parkash, Rajpurohit and Ramniwas, 2008; Hymenoptera, Formicidae - Kaspari *et al.*, 2015) with closely related species demonstrating significantly different thermal tolerances. The extension of termite physiological tolerances may have enabled them to withstand the more variable conditions and colonise savannas. However, few studies have documented the physiological limits of soil insects, including termites (but see, Woon et al., 2018), so the patterns observed in other studies may not hold across all insect orders.

In recent years, an increased focus on the thermal niche of species (Gvoždík, 2018; Kearney et al., 2010; Porter and Kearney, 2009; Sunday et al., 2014), has produced hypotheses to explain thermal performance, particularly given climate change concerns (Buckley et al., 2013; Clusella-Trullas et al., 2011; Huey et al., 2012; Kearney and Porter, 2009; Sunday et al., 2011). These studies and hypotheses have contributed to developing the Thermal Adaptation Hypothesis (Angilletta et al., 2002; Angilletta, 2009; Huey and Kingsolver, 1989; Kaspari et al., 2015; Kingsolver and Gomulkiewicz, 2003), which states that species should be adapted to the thermal niche of their environment, therefore suggesting that species found in thermally more variable environments should have wider thermal ranges (Angilletta, 2009). In addition, the thermal adaptation hypothesis suggests that species in more thermally stable environments, as well as having narrower thermal ranges, should be operating closer to their thermal optimum (Angilletta, 2009). In the context of our study system, tropical forests have much more stable environmental conditions, whereas savannas have more extreme limits and more variation, so we would expect savanna species to have wider and more extreme physiological limits to adapt to their environment.

Here, we test the hypothesis that savanna termite species have wider physiological limits, which are likely to have facilitated their colonisation and speciation within savanna ecosystems. We quantify the thermal limits (critical thermal minima [CT_min_] and critical thermal maxima [CT_max_]) of the most abundant mound-building termite species across tropical forest and savanna areas in Ghana, to test whether savanna species have more extreme physiological limits than forest species. We assess these data and discuss the applicability of the thermal adaptation hypothesis.

We predict from our hypothesis that (1) termite species from savannas will have higher CT_max_ values than forest species, and that (2) they will have lower CT_min_ values. We correlated the physiological tolerances with climatic factors that change across the environmental gradient (from forest to savanna), temperature, rainfall and canopy cover, to determine which, if any, may be more strongly associated with physiological limits.

## Materials and Methods

### Sampling Locations

Termites were sampled from four locations in Ghana across two different biomes: tropical rainforest and Guinea savanna. The three forest locations were: Bobiri Forest Reserve, a moist semi-deciduous forest in the Ashanti Region, a matrix of forest patches that have been, or currently are, logged (all colonies sampled from fragments that have not been logged for at least 20 years); the Forestry Research Institute of Ghana (FoRIG) Campus, a moist semi-deciduous forest patch located on the campus at Fumesua, near Kumasi, in the Ashanti Region; and the Assin-Attandanso Resource Reserve side of the Kakum Conservation Area, a moist evergreen tropical forest that has been unlogged since 1990 (Wiafe, 2016), in the Central Region. Termites were sampled from a single savanna site, Mole National Park, a large unfenced protected area in the Savannah Region, between June and July 2019. Sampling in all locations was conducted during the end of the dry season and the start of the wet season. Termites were sampled during the end of the dry season and start of the wet season at all sites: between January and March 2019 for the three forest sites, and June and July 2019 for the savanna site (Table 1).

**Table 1.** Summary statistics of the different sampling sites, with their locations and climatic conditions. The GPS co-ordinates are a representative location, at the approximate centre of the sampling location. Climate data sourced from WorldClim 2 (Fick and Hijmans, 2017).

| Location | Vegetation type | GPS Co-ordinates | Mean Maximum<br>Temperature | Mean Minimum<br>Temperature | Mean Annual<br>Rainfall |
| --- | --- | --- | --- | --- | --- |
| Bobiri Forest Reserve | Rainforest | 6.689 N,<br>1.342 W | 31.3°C | 18.5°C | 1306 mm |
| Forestry Research Institute<br>of Ghana (FoRIG) | Rainforest | 6.716 N,<br>1.529 W | 31.4°C | 19°C | 1357 mm |
| Kakum Farmland | Farmland (formerly forest) | 5.568 N,<br>1.388 W | 31°C | 19.8°C | 1504 mm |
| Kakum National Park | Rainforest | 5.573 N,<br>1.379 W | 30.9°C | 19.7°C | 1502 mm |
| Mole North | Savanna | 9.334 N,<br>1.868 W | 35.2°C | 19.3°C | 1018 mm |
| Mole South | Savanna | 9.286 N,<br>1.847 W | 35.2°C | 19.3°C | 1025 mm |

### Termite and Environmental Sampling

We sampled termites directly from termite mounds, identified to genus level (based on their diagnostic mound structures), which we located using directed searching, using the knowledge and experience of the field assistants. Two mounds were sampled per day, typically between 07h00 and 09h00. Target mounds were broken open and we collected termites, as well as some mound material, and placed them into a cool box to protect the termites from external conditions and reduce stress that could affect their thermal tolerances while being transported to the laboratory. The sample from each colony was placed in a separate container to prevent interaction between the two colonies. Where possible, both worker and soldier castes were sampled. The time between sampling and experimentation was reduced as much as possible (mean = 122.6 mins, sd = ±81.4 min). The large standard deviation is attributed to having a single collection, but running two experiments, per day. The same colonies that were sampled were used in two experiments, one testing thermal maximum and one testing thermal minimum (the order of which was decided randomly); hence, the individuals used in the second experiment were kept inside the cool box (which was out of direct sunlight) while the first experiment took place.

We photographed each mound to assist with species identification. Canopy cover above each sampled mound was quantified using a hemispherical fish-eye lens (Canon EOS 70D camera, Sigma 4.5mm f/1:2.8 Circular Fisheye Lens). For large mounds the image was taken directly above the apex of the mound, but with small mounds (<75 cm in height) the image was taken above the mound at a height of 75 cm due to the height of the stabilising monopod. Due to variable light conditions when the images were taken, we did not standardise ISO, shutter-speed or aperture, although variation in these settings were reduced as much as possible. Variable light conditions and camera settings were corrected using Gap Light Analyzer software (GLA; Frazer et al., 1999). The approximate leaf-area index above each mound was then calculated as a measure of canopy cover. Leaf-area index is a measurement of the quantity of leaf area in a canopy per unit area of ground, and ranges from 0 (no leaf cover) to 6 (complete leaf cover).

### Thermal Tolerance Experiments

We used similar methods to those outlined in Bishop et al. (2017) to estimate the thermal tolerance of the termites. Fourteen individuals from each of two colonies (28 in total) were tested per experimental run. Only individuals of the helper castes, workers and soldiers, were used in the experiments Each termite was placed into a separate microcentrifuge tube. The tubes were then placed into a dry heat bath (Tropicooler 260014-2, Boekel Scientific, Feasterville, PA, USA; temperature range −19°C to 69°C and an accuracy of ±1°C). The heat bath contained two aluminium inserts with each consisting of 14 wells, each of which held a single microcentrifuge tube. Individuals from two colonies and different castes were randomly allocated to different wells to remove potential equipment bias. The two colonies sampled each day were used in each experimental run (CT_max_ and CT_min_) and each colony was tested for both CT_max_ and CT_min_; however, each individual termite was only used in one experiment.

The temperature within each microcentrifuge tube was acclimated for 15 minutes. CT_max_ experiments had an acclimation temperature of 30°C, and CT_min_had an acclimation temperature of 24°C. After the acclimation period, the temperature was changed by 1°C (raised for a CT_max_ experiment, and lowered for CT_min_), and maintained at each new temperature for a further three minutes. Following this three-minute period, we checked each termite for loss of muscle co-ordination (CT_max_) or lack of movement (CT_min_) by flicking the microcentrifuge tube. If we noticed the termite lost muscle co-ordination (CT_max_) or had stopped moving (CT_min_) the temperature was taken as the thermal limit of that individual termite. The experimental run was ended when the thermal limit of all 28 termites was recorded.

### Termite Identification

Termite colonies were identified to genus level in the field based on their mound structure. All termites used in experimental runs were preserved in 70% ethanol and brought to the UK. We took an individual from each colony and DNA was extracted from each sample. The *COII* mitochondrial gene was amplified and sequenced using Sanger Sequencing at the Natural History Museum, London. Each sequence was matched with the most closely related sequence on NCBI using BLAST (Johnson et al., 2008). If the sequences had a unique high likelihood identity match (>98%) with a single species described on the NCBI database, that sample, and therefore the colony it was sampled from, was considered identified as that species. If there was not a unique high likelihood identity match, the sequences (and therefore the colonies from which each sample was taken) were grouped and labelled as an unspecified species within a genus (for example, *Cubitermes sp. A*, Fig. 1). After identification, species that did not have at least three replicates per sampling location were removed from the dataset (19 colonies removed, present in online dataset). Since the genetic identification was conducted, the genus *Cubitermes* was split into several genera (Hellemans et al., 2020). However, there were limited sequence data for the new classifications so we have treated all the new genera as *Cubitermes sensu lato*.

**Fig. 1.**
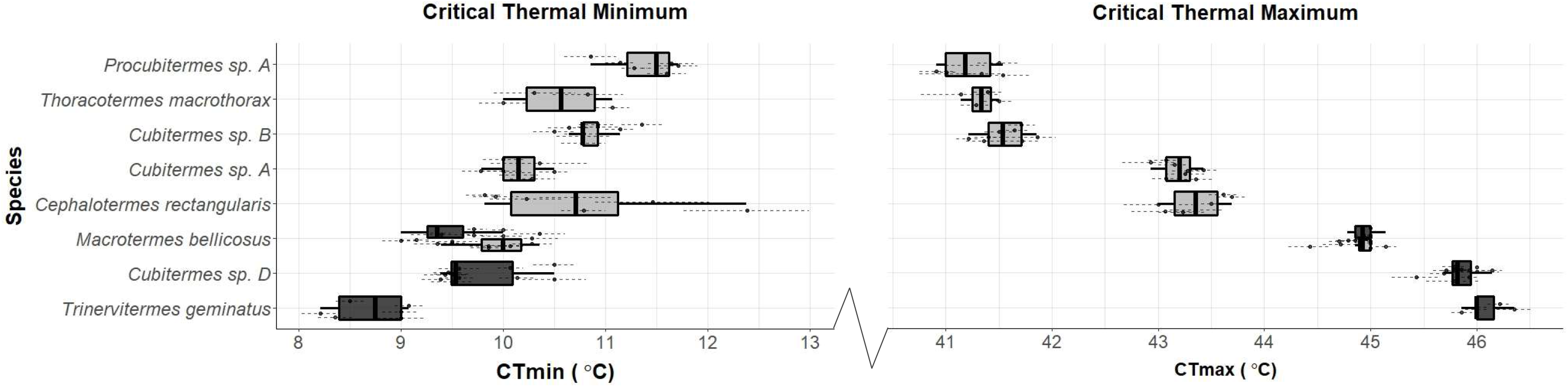
Relationship between termite species thermal limits and the habitat they were sampled from. Species are ordered from lowest CTmax (top) to highest CTmax (bottom). Colour of the boxplot denotes the location the termites were sampled, dark-grey from savanna and light-grey from forest. Macrotermes bellicosus was sampled from both savanna and forest environments, and so it has distinct boxplots for each community. The points represent the estimated thermal limit of a unique termite colony, and the dotted lines represent the standard error around the colony average. Central thick line of the boxplot represents the species median thermal limit, the box represents the interquartile range, and the whiskers the upper and lower adjacent values.

### Climate Data

Climate data was sourced from the WorldClim2 dataset (Fick and Hijmans, 2017) using the ‘raster’ package (Hijmans et al., 2020) in the R statistical software version 3.6.2 (R Core Team, 2019). WorldClim2 gets average measurements of climatic variables each month, based on an interpolated dataset between 1970 and 2000. We extracted measurements of average monthly rainfall, average monthly maximum temperature, and average monthly minimum temperature to the highest resolution possible (30 arc seconds, equivalent to approximately 1km^2^) for each mound we sampled. From these data we obtained average daily rainfall, highest average maximum temperature and lowest average minimum temperature for the area around each colony centre.

## Statistical Analysis

After genetic identification, eight species had a sample size of three colonies and were included in the statistical analyses. Three of those species were sampled from the two savanna locations (Mole North, Mole South, Table 1), and six species were sampled from the forest locations (FoRIG, Bobiri, Kakum Forest and Kakum Farmland, Table 1). One species, *Macrotermes bellicosus*, was sampled in both savanna and forest locations, but it was only found in the forest environment that was heavily disturbed, Kakum Farmland (Table 1).

All statistical analyses used the R statistical package version 3.6.2 (R Core Team, 2019). We calculated the estimated colony CT_max_ and CT_min_ by taking the average of the thermal tolerance of each termite within that colony that was used in an experimental run. Initially, basic Welch’s two sample t-tests were run to compare the thermal tolerances of species assemblages sampled in forest and savanna environments (termites sampled from Kakum Farmland were not included, Table 1). A t-test was also used to compare *Macrotermes bellicosus* thermal tolerances in two different environment types (farmland and savanna), and a one-way analysis of variance test was used to compare the thermal tolerances of the three *Cubitermes* species sampled. Tukey’s HSD post-hoc statistics were used for pairwise comparisons of the species thermal tolerances.

To test whether environmental variables affect termite colony thermal limits, mixed effects models were fitted using the R package ‘lme4’ (Bates et al., 2019), with a separate model for CT_max_ and CT_min_. Maximal models tested the interactive effects of canopy cover, average daily rainfall (which was rescaled from annual rainfall) at the mound, and temperature at the mound (maximum temperature for the CT_max_ model, and minimum temperature for the CT_min_ model). A single random effect of colony location (as listed in Table 1, Fig. S1) was used to account for additional unmeasured environmental differences. Species was not included as a random effect (crossed or nested) because there was little overlap in species composition between sampling locations, and so both site and species would therefore explain the same variation, and so there was not enough data to prevent non-convergence.

To understand which of the fixed effects and their interactions best explained the variation in termite thermal tolerances, two lists of candidate models were generated (one for CT_max_ and one for CT_min_) using the ‘dredge’ function from the ‘MuMIn’ package (Bartoń, 2019). This takes the maximal model and systematically removes variables and/ or interactions until all possible combinations have been tested, and compared these complete lists of models using the package ‘AICcmodavg’ (Mazerolle and Linden, 2019). The model with the lowest corrected AIC (AICc) value was obtained, and the other models with a difference in AICc (ΔAICc) of less than two were retained to inform the final ‘full’ model (Table S1 and Table S2; Burnham and Anderson, 2002). Model averaging, using the candidate lists of the model with the lowest AICc value and all those with ΔAICc < 2, was used to identify the best predictor variables across the models. The relative importance of these predictors was calculated as the sum of their relative Akaike weights from all models in which they appear; thus, models with lower AICc values contribute more to the final ‘full’ model. A series of *F* tests was also conducted to verify the significance of the retained parameters on the thermal limits of the termite species tested in the final models. This process was conducted for two separate model sets, one testing CT_min_ and the other CT_max_, generating two ‘full’ models.

## Results

A total of 2042 termite specimens from 71 colonies and eight species were used in the final analyses (1022 termites for CT_max_ experiments, 1020 termites for CT_min_ experiments). There was a large amount of interspecific variation in thermal tolerance, with species averages varying by 4.86°C for their CT_max_ and 2.96°C for their CT_min_ (Fig. 1). Savanna species had significantly higher CT_max_ values on average than forest species (p < 0.001, t = −17.309, df = 52.37, Fig. 1A), and significantly lower CT_min_ values on average than forest species (p < 0.001, t = 8.7035, df = 53.84, Fig. 1B); as a result, savanna species had wider thermal limits than forest species. The three *Cubitermes* species tested had significantly different CT_max_ values (p < 0.001, F_2,26_ = 1424, Fig. 1A) and CT_min_ values (p < 0.001, F_2,26_ = 29.24, Fig. 1B), with all species having significantly different thermal niches from one another (Tukey’s HSD; Fig. S2 & Fig. S3). *Macrotermes bellicosus* from multiple environments was tested (farmland in the forest zone and savanna), but CT_max_ did not differ among the two habitats (p = 0.805, t = 0.255, df = 7.775, Fig. 1A). *M. bellicosus* did, however, have significantly different CT_min_ values (p = 0.008, t = 3.132, df = 12.266, Fig. 1B), with savanna colonies having lower CT_min_.

## Critical thermal maximum

The full model retained all three fixed effects (canopy cover, maximum temperature and average daily rainfall; Table 2), but the interaction between the terms were removed during the model selection process. Of the three remaining variables, only canopy cover had a significant negative correlation with colony CT_max_ (z = 3.138, p = 0.001, df = 63.470; Fig. 1A; Table 2), suggesting that as the canopy cover above termite mounds increases, the average critical thermal maximum of the termite colony decreases (Fig. 2). Despite being retained in the full model, neither maximum temperature (z = 1.215, p = 0.358, df = 3.900; Table 2) nor rainfall (z = 0.068, p = 0.964, df = 3.689; Table 2) showed a significant correlation with colony CT_max_. The single random effect of colony location (Table 1) explained 48.6% of the variation in the model.

**Table 2.**
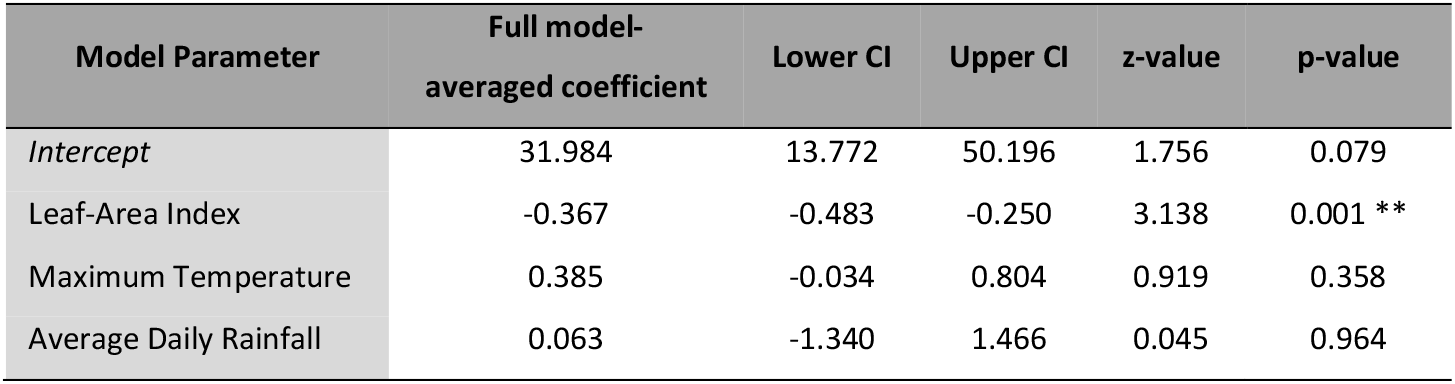
‘Full’ averaged model coefficients and their significance as produced by the ‘AICcmodavg’ package (Mazerolle and Linden, 2019) modelling termite CT_max_ change as a function of environmental factors. The full model fitted the following fixed effects and a single random effect of sampling location to predict the critical thermal maximum of termite colonies. Asterisks indicate level of significance.

**Fig. 2.**
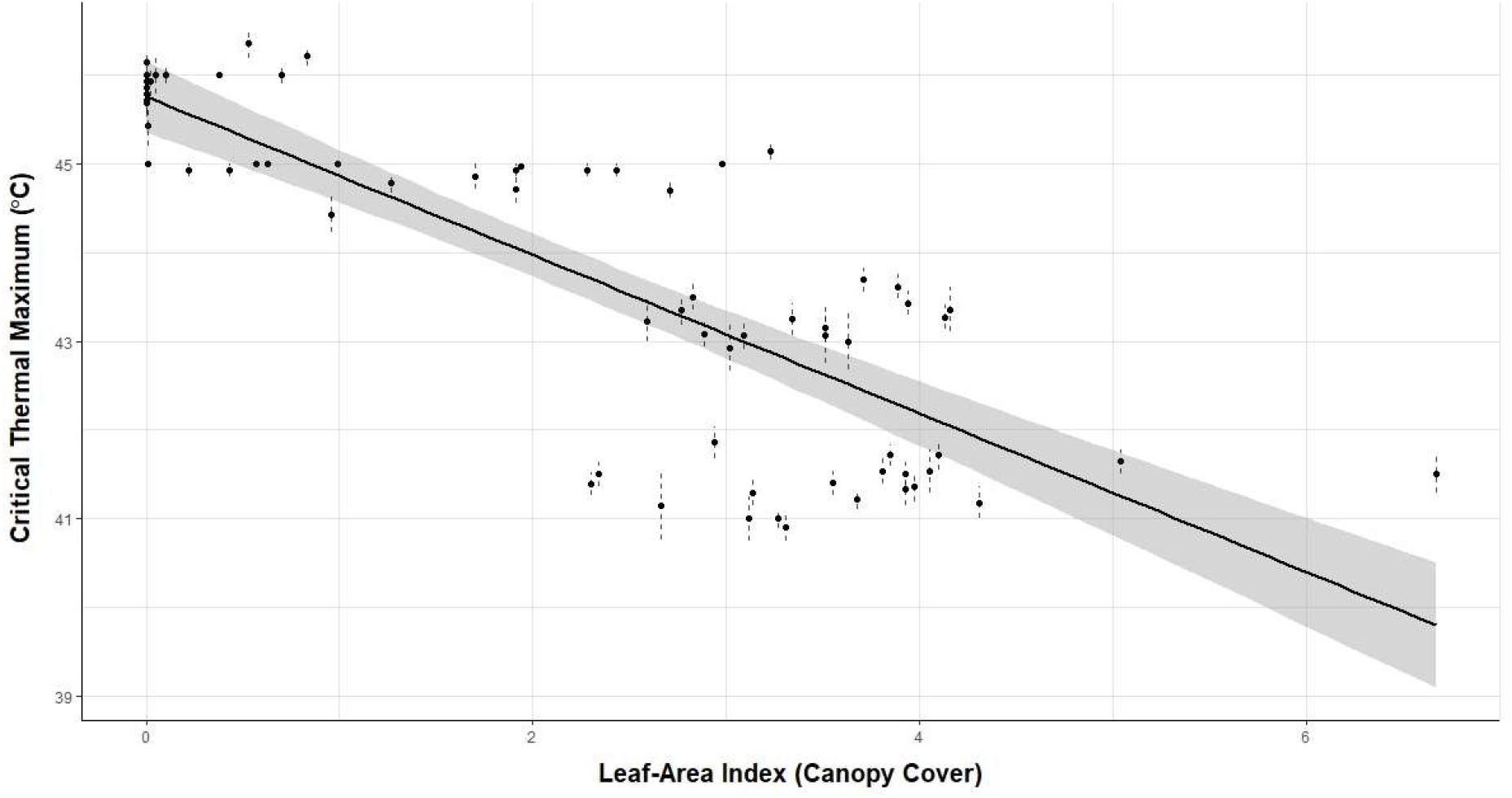
Significant negative correlation between the estimated critical thermal maximum of termite colonies and the canopy cover above the colony centre (p = 0.001), as predicted by a linear mixed effects model (with sampling location as the random effect). Leaf-area index is the measurement of canopy cover used, with higher leaf-area index representing higher canopy cover above the mound. Each point represents the estimated thermal maximum of a unique termite colony across all locations sampled, the thermal maximum is averaged from each individual termite that was used in the experiment, and the dotted line represents the standard error around this mean. The solid black line represents predicted relationship based on the model, with the shaded grey area representing the uncertainty around these predictions.

**Fig. 3.**
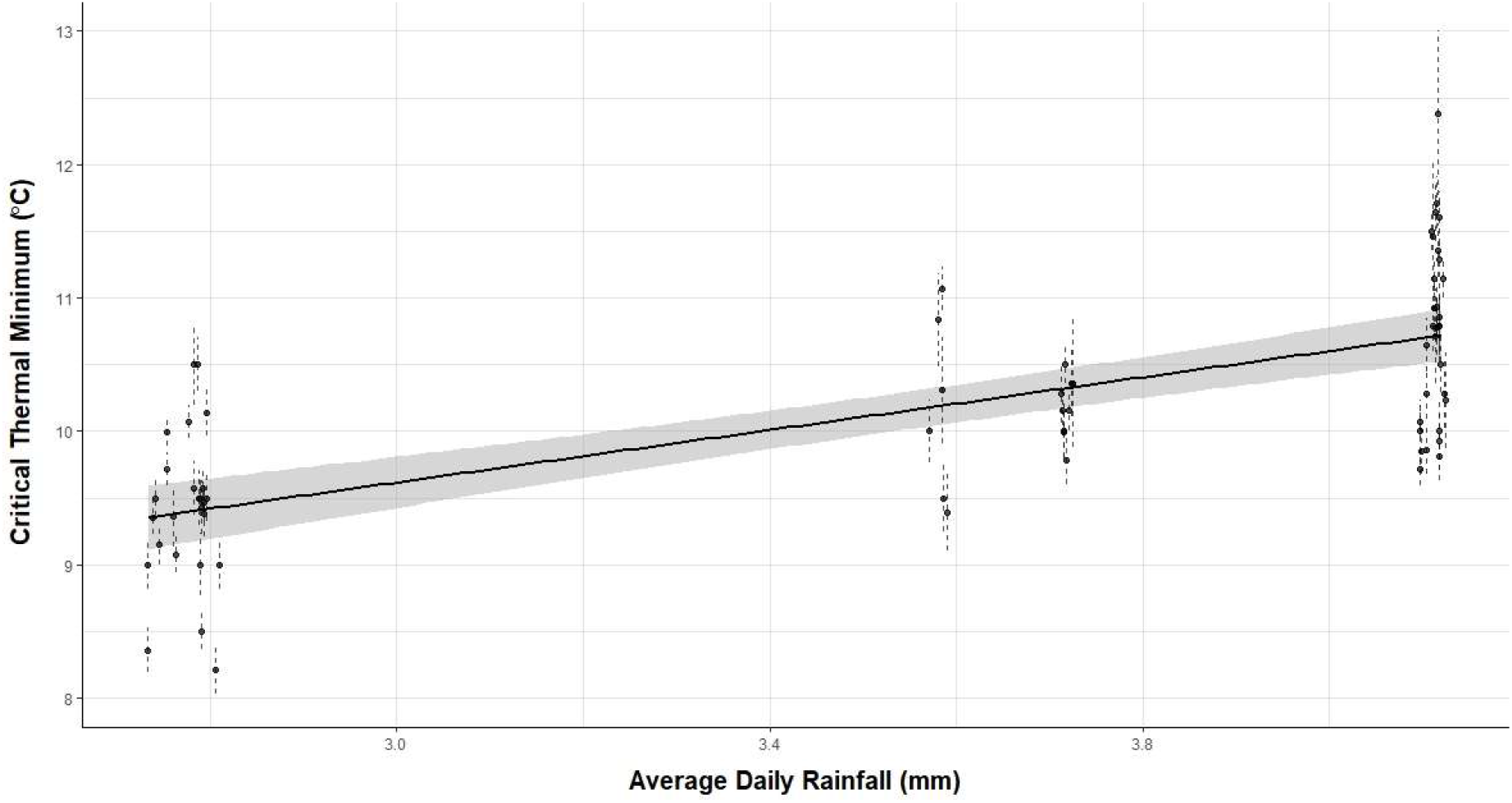
Significant positive correlation between average daily rainfall at the mound (data obtained from WorldClim2, Fick and Hijmans, 2017), and the estimated critical thermal minimum of termite colonies (p = 0.007), as predicted by a linear mixed effects model (with sampling location as the random effect). Each point represents the estimated thermal minimum of a unique termite colony across all locations sampled, the thermal minimum is averaged from each individual termite that was used in the experiment, and the dotted line represents the standard error around this mean. The solid black line represents predicted relationship based on the model, with the shaded grey area representing the uncertainty around these predictions.

## Critical thermal minimum

The full critical thermal minimum model retained two fixed effects (canopy cover and average daily rainfall; Table 3), and it also removed all interactions between the variables. Unlike in the CT_max_ model, canopy cover was not significantly correlated with CT_min_ of termite colonies (z = 0.269, p = 0.788, df = 59.671; Fig. 1B; Table 3); however, average daily rainfall was significantly positively correlated with CT_min_ (z = 2.679, p = 0.007, df = 4.504; Fig. 1B; Table 3). Colony location (Table 1) explained 39.6% of variation in the model.

**Table 3.** ‘Full’ averaged model coefficients and their significance as produced by the ‘AICcmodavg’ package (Mazerolle and Linden, 2019) modelling termite CT_min_ change as a function of environmental factors. The full model fitted the following fixed effects and a single random effect of sampling location to predict the critical thermal minimum of termite colonies. Asterisks indicate level of significance.

| Model Parameter | Full model-<br>averaged coefficient | Lower CI | Upper CI | z-value | p-value |
| --- | --- | --- | --- | --- | --- |
| Intercept | 6.878 | 5.714 | 8.042 | 5.910 | 2e-16 *** |
| Average Daily Rainfall | 0.885 | 0.555 | 1.216 | 2.679 | 0.007 ** |
| Leaf-Area Index | 0.011 | -0.030 | 0.052 | 0.269 | 0.788 |

## Discussion

Our findings support the hypothesis that savanna termite species have more extreme physiological tolerances, and developing these tolerances is likely to have been a key mechanism that has allowed termites to colonise savanna environments. As predicted, savanna species had a significantly higher CT_max_ than forest species. In addition, CT_max_ had a significant negative correlation with canopy cover. As the canopy cover decreases the amount of sunlight reaching the soil surface and the surface of the mound increases, which will increase the temperature of the air, soil surface, and potentially within the mound and foraging areas (Hardwick et al., 2015). Canopy cover in our savanna environments (average 26.6 ± 5.8%) was much lower than that found in tropical forests (84.7 ± 3.1%). To cope with higher day temperatures in savannas, the termites will have had to develop mechanisms such as increased thermal tolerance. Our data support this idea, not only due to the positive correlation between canopy openness and CT_max_, but also among closely related species (e.g., of *Cubitermes*) in which species in the more open and extreme savanna habitat had more extreme thermal limits (Fig. 1A & Fig. 1B). *Cubitermes* are soil feeding termites, so do not need to leave the protection of the soil to forage for food and so have a reduced interaction with ambient conditions (when compared to other species), yet they have significantly different thermal tolerances between species from the two environments. If these species had developed extensive mound thermoregulatory mechanisms such that the environmental conditions experienced by the termites were similar, we would expect closely related (i.e., congeneric) species to have similar thermal tolerances, irrespective of the location they were sampled from; this is not the case with *Cubitermes*, as closely related species have significantly different thermal tolerances.

Surprisingly, despite savanna species having a significantly lower CT_min_ than forest species, CT_min_ was not significantly correlated with canopy cover. We expected that lower canopy cover in savannas would allow the ground surface to cool more rapidly at night, and result in lower temperatures. Consequently, we expected canopy cover to positively correlate with CT_min_, as termites in locations with lower canopy cover will experience lower temperatures at night. This is not what was found; CT_min_ was significantly positively correlated with average daily rainfall. We are unsure about the underlying process linking lower CT_min_ with reduced rainfall. The species from the savanna regions had lower CT_min_ values, which has lower rainfall and therefore lower cloud cover, which would increase heat loss and result in lower temperatures; however, minimum temperature was included in the model and had no significant effect. More sampling is required, additional species from more locations along the environmental gradient, including between the forest and savanna environments, and further north of our savanna sites (in locations which would have higher day temperatures, and lower night temperatures) to properly tease apart these relationships.

Canopy cover was expected to have a significant relationship with CT_min_ because some termite species forage during cool parts of the day, including night time, which would put pressure on the CT_min_ of foraging individuals, and that pressure would be greater in areas with low canopy cover where the less heat would be retained overnight. For example, the specialist savanna grass-feeder, *Trinervitermes*, typically forages at dusk, dawn or during the night (Adam et al., 2008). Our data support this: *Trinervitermes germinatus* had the lowest CT_min_, which would be expected for coping with the lower temperatures experienced during foraging at night. Other studies have shown that some savanna species (e.g. *Macrotermes* sp.; Darlington, 1982) forage during cooler parts of the day or when overcast (Evans and Gleeson, 2001; Parr pers obs); however, these studies are few in number and do not cover the species studied here. In general, behavioural avoidance of one thermal extreme (e.g., in the case of *Trinervitermes*, hot conditions) may reduce selection pressures on the physiological tolerance of that extreme (e.g. CT_max_), but not necessarily the opposite extreme. This interplay between behavioural and physiological adaptations represents an important avenue for future study.

Mean site temperature did not have a significant effect on thermal tolerance at maximum or minimum temperatures. This may be because the temperature data used in both models was interpolated temperature of air (Fick and Hijmans, 2017), which is unlikely to be the best measure of the conditions that termites would experience. Due to their small size, terrestrially foraging termites forage in the thermal boundary layer, immediately above a surface of, in this case, the ground. The thermal boundary layer has very different conditions from the recorded air temperature, both during the day (where it is likely to be super-heated) and at night (retaining more heat due to proximity to soil). Boundary layer temperature, or ground surface temperature, would be a more accurate representation of the true thermal conditions that termites experience, but unfortunately these data were not available at a fine-enough spatial scale. So, while it is likely that air temperature and ground temperature are correlated, and lack of ground temperature does not invalidate these results, the model would be improved by access to these data.

There was more intraspecific variation in CT_min_ than CT_max_ possibly associated with measurement error. The protocol for recording CT_min_ used for other insects, such as ants (Bishop et al., 2018; Kaspari et al., 2015), involves recording the temperature at which all movement ceases. However, termites are much less mobile than ants, making it extremely difficult to tell the difference between the onset of a chill coma (Hazell and Bale, 2011) and the true CT_min_. Therefore, whereas the values described here are biologically relevant, as termite fitness would be severely negatively affected if temperatures were to drop to near these recorded “CT_min_” values, the values recorded likely contain measurement error. An additional method to test for signs of life in other insects is to shine a laser pen into the eye of the individual in an attempt to elicit a head-movement response. This method is not possible with termites, as the helper castes of almost all species lack eyes, so the only stimulus that could be used was flicking the test tube. Another source of error is that a ‘no movement’ reaction in a living individual is impossible to distinguish visually from a dead individual. Consequently, these measurements may slightly underestimate true CT_min_, resulting in greater intraspecific variation than the true range. However, the measurement errors are likely to be random, rather than systematically correlated with environmental variables. Moreover, despite the measurement inaccuracies, CT_min_ correlated significantly with rainfall, and the differences in CT_min_ between the species were pronounced.

Although we think it unlikely that there are mound thermoregulatory mechanisms in the *Cubitermes* species we sampled because of their very different thermal limits, our data are equivocal as to whether there is mound thermoregulation in *Macrotermes bellicosus* mounds. This is a topic of much debate (Korb, 2003; Jones and Oldroyd, 2006; Gouttefarde et al., 2017; Scott Turner, 1994; Turner, 2001). *Macrotermes bellicosus* had the same CT_max_ in all locations sampled despite environmental conditions varying among habitats; this suggests that there may be thermoregulatory mechanisms within their mounds, likely to maintain ideal growing conditions for their fungus symbionts (Korb and Linsenmair, 2000; Korb, 2003). We sampled them from their native region (savanna) and further south of their ancestral distribution, areas which have less extreme environmental conditions, particularly a much lower maximum temperature, which would not put any additional pressure on their thermal limits. Improved knowledge of likely thermoregulation in *Macrotermes* mounds could come from comparisons of thermal tolerances of *M. bellicosus* from multiple savanna environments, with varying climatic conditions (e.g. further north from where we sampled, with a hotter and drier climate).

As predicted by the thermal adaptation hypothesis, the forest species had much narrower thermal limits than the savanna species (Angilletta, 2009). Forests have more stable temperatures (less diurnal and annual variation), so the theory posits that the termite species found there should have narrower thermal limits than the savanna species. This prediction of the thermal adaptation hypothesis holds true, and it is likely that the stability, and lower extremes, of forest environments have contributed to narrower thermal limits (and therefore thermal niches) of forest termite species. How close the current temperatures are to termite thermal limits is not known, however, due to the imprecise nature of the temperature data. Obtaining more accurate temperate data, which measures surface temperature or boundary layer temperature, over a finer spatial scale, and recorded from a more recent time period would allow us to make a more accurate prediction of the resilience of termite assemblages to a warming climate. Understanding how close the thermal conditions that termites experience are to their thermal limits, and to thermal optima, is vital in making this prediction.

Our results suggest that for termites, the most important soil macroinvertebrates in savanna environments (Davies et al., 2016a; Jouquet et al., 2016), a key mechanism that has facilitated their colonisation of the seasonal, climatically extreme environments of savannas from forests is the widening of their thermal tolerances. However, temperature is not the only variable that changes along an environmental gradient from forest to savanna. Food type, food availability, soil mineral content (and pH), humidity, predator diversity and abundance are a few of the factors that could change along the gradient, and therefore influence termite species distributions. Some of these would contribute to the large proportion of variation that is explained by the random effect of sampling location in both models, making them promising avenues for future research. For example, we have shown previously that desiccation tolerance is low in Bornean forest termites (Woon et al., 2018), so quantifying the desiccation tolerance of African species, and comparing these tolerances between environment types could provide additional information on how certain species are able to survive in savannas.

Understanding how termite physiology and behaviour changes across environmental gradients such as this one is particularly important in the context of climate change. The thermal adaptation hypothesis predicts that species in more stable environments will be more heavily impacted by warming conditions, due to those species operating closer to their thermal maximum (Angilletta, 2009), and be more susceptible to extinction. This suggests the forest species here may be more vulnerable to rising temperatures associated with climate change, but more research is needed to understand the complexities of physiological change, how it affects termite distributions, and how they might be altered by the changing climate.

Our data suggest that the widening of termite thermal tolerances has occurred in species that have established in areas of more extreme climatic conditions, and that is likely a large factor in allowing termites to persist, speciate, and become ecologically dominant in arid, tropical environments. Whether the widening of these physiological limits will make them more resilient to climate change is a question that remains to be answered.

## References

Aanen, D. K. and Eggleton, P. (2005). Fungus-growing termites originated in African rain forest. Current Biology 15, 851–855.

Adam, R. A., Mitchell, J. D. and Westhuizen, M. C. van der (2008). Aspects of foraging in the harvester termite, *Trinervitermes trinervoides* (Sjöstedt) (Termitidae: Nasutitermitinae). African Entomology 16, 153–161.

Adjorlolo, C., Mutanga, O., Cho, M. A. and Ismail, R. (2012). Challenges and opportunities in the use of remote sensing for C3 and C4 grass species discrimination and mapping. African Journal of Range & Forage Science 29, 47–61.

Angilletta, M. J. (2009). Thermal Adaptation: A Theoretical and Empirical Synthesis. Oxford University Press.

Angilletta, M. J., Niewiarowski, P. H. and Navas, C. A. (2002). The evolution of thermal physiology in ectotherms. Journal of Thermal Biology 27, 249–268.

Bartoń, K. (2019). MuMIn: Multi-Model Inference. https://CRAN.R-project.org/package=MuMIn

Bates, D., Maechler, M., Bolker, Walker S., Christensen, R. H. B., Singmann, H., Dai, B., Scheipl, F., Grothendieck, G., Green, P., & Fox, J. (2019). lme4: Linear Mixed-Effects Models using “Eigen” and S4. https://CRAN.R-project.org/package=lme4

Bishop, T. R., Robertson, M. P., Van Rensburg, B. J. and Parr, C. L. (2018). Coping with the cold: minimum temperatures and thermal tolerances dominate the ecology of mountain ants. Ecological Entomology 105–114.

Bottinelli, N., Jouquet, P., Capowiez, Y., Podwojewski, P., Grimaldi, M. and Peng, X. (2015). Why is the influence of soil macrofauna on soil structure only considered by soil ecologists? Soil and Tillage Research 146, 118–124.

Bouchenak-Khelladi, Y., Slingsby, J. A., Verboom, G. A. and Bond, W. J. (2014). Diversification of C4 grasses (Poaceae) does not coincide with their ecological dominance. American Journal of Botany 101, 300–307.

Bourguignon, T., Lo, N., Cameron, S. L., Šobotník, J., Hayashi, Y., Shigenobu, S., Watanabe, D., Roisin, Y., Miura, T. and Evans, T. A. (2015). The Evolutionary History of Termites as Inferred from 66 Mitochondrial Genomes. Molecular Biology & Evolution 32, 406–421.

Buckley, L. B., Tewksbury, J. J. and Deutsch, C. A. (2013). Can terrestrial ectotherms escape the heat of climate change by moving? Proceedings of the Royal Society B: Biological Sciences 280, 20131149.

Burnham, K. P. and Anderson, D. R. (2002). Model selection and multimodel inference: A practical information-theoretic approach. In A Primer on Natural Resource Science, p. Texas A&M University Press.

Cerling, T. E., Wang, Y. and Quade, J. (1993). Expansion of C4 ecosystems as an indicator of global ecological change in the late Miocene. Nature 361, 344–345.

Cerling, T. E., Harris, J. M., MacFadden, B. J., Leakey, M. G., Quade, J., Eisenmann, V. and Ehleringer, J. R. (1997). Global vegetation change through the Miocene/Pliocene boundary. Nature 389, 153–158.

Christin, P.-A., Osborne, C. P., Chatelet, D. S., Columbus, J. T., Besnard, G., Hodkinson, T. R., Garrison, L. M., Vorontsova, M. S. and Edwards, E. J. (2013). Anatomical enablers and the evolution of C4 photosynthesis in grasses. PNAS 110, 1381–1386.

Clusella-Trullas, S., Blackburn, T. M. and Chown, S. L. (2011). Climatic predictors of temperature performance curve parameters in ectotherms imply complex responses to climate change. The American Naturalist 177, 738–751.

Dangerfield, J. M., Mccarthy, T. S. and Ellery, W. N. (1998). The mound-building termite *Macrotermes michaelseni* as an ecosystem engineer. Journal of Tropical Ecology 14, 507– 520.

Darlington, J. P. E. C. (1982). The underground passages and storage pits used in foraging by a nest of the termite *Macrotermes michaelseni* in Kajiado, Kenya. Journal of Zoology 198, 237–247.

Davies, A. B., Robertson, M. P., Levick, S. R., Asner, G. P., Rensburg, B. J. van and Parr, C. L. (2014). Variable effects of termite mounds on African savanna grass communities across a rainfall gradient. Journal of Vegetation Science 25, 1405–1416.

Davies, A. B., Levick, S. R., Robertson, M. P., Rensburg, B. J. van, Asner, G. P. and Parr, C. L. (2016a). Termite mounds differ in their importance for herbivores across savanna types, seasons and spatial scales. Oikos 125, 726–734.

Davies, A. B., Baldeck, C. A. and Asner, G. P. (2016b). Termite mounds alter the spatial distribution of African savanna tree species. Journal of Biogeography 43, 301–313.

Davies, T. J., Daru, B. H., Bezeng, B. S., Charles-Dominique, T., Hempson, G. P., Kabongo, R. M., Maurin, O., Muasya, A. M., van der Bank, M. and Bond, W. J. (2020). Savanna tree evolutionary ages inform the reconstruction of the paleoenvironment of our hominin ancestors. Scientific Reports 10, 12430.

Dierks, A. and Fischer, K. (2008). Feeding responses and food preferences in the tropical, fruit-feeding butterfly, *Bicyclus anynana*. Journal of Insect Physiology 54, 1363–1370.

D’Onofrio, D., Sweeney, L., von Hardenberg, J. and Baudena, M. (2019). Grass and tree cover responses to intra-seasonal rainfall variability vary along a rainfall gradient in African tropical grassy biomes. Scientific Reports 9, 2334.

Eggleton, P. (2000). Global patterns of termite diversity. In Termites: Evolution, Sociality, Symbioses, Ecology (ed. Abe, T., Bignell, D. E., and Higashi, M.), pp. 25–51. Dordrecht: Springer Netherlands.

Evans, T. A. and Gleeson, P. V. (2001). Seasonal and daily activity patterns of subterranean, wood-eating termite foragers. Australian Journal of Zoology 49, 311–321.

Fick, S. E. and Hijmans, R. J. (2017). WorldClim 2: new 1-km spatial resolution climate surfaces for global land areas. International Journal of Climatology 37, 4302–4315.

Fox-Dobbs, K., Doak, D. F., Brody, A. K. and Palmer, T. M. (2010). Termites create spatial structure and govern ecosystem function by affecting N2 fixation in an East African savanna. Ecology 91, 1296–1307.

Frazer, G. W., Canham, C. D. and Lertzman, K. P. (1999). Gap Light Analyzer (GLA), Version 2.0: Imaging software to extract canopy structure and gap light transmission indices from true-colour fisheye photographs.

Gouttefarde, R., Bon, R., Fourcassié, V., Arrufat, P., Haifig, I., Baehr, C. and Jost, C. (2017). Investigating termite nest thermodynamics using a quick-look method and the heat equation. bioRxiv 161075.

Gvoždík, L. (2018). Just what is the thermal niche? Oikos 127, 1701–1710.

Halali, S., Brakefield, P. M., Collins, S. C. and Brattström, O. (2020). To mate, or not to mate: The evolution of reproductive diapause facilitates insect radiation into African savannahs in the Late Miocene. Journal of Animal Ecology 89, 1230–1241.

Hardwick, S. R., Toumi, R., Pfeifer, M., Turner, E. C., Nilus, R. and Ewers, R. M. (2015). The relationship between leaf area index and microclimate in tropical forest and oil palm plantation: Forest disturbance drives changes in microclimate. Agricultural and Forest Meteorology 201, 187–195.

Hazell, S. P. and Bale, J. S. (2011). Low temperature thresholds: Are chill coma and CTmin synonymous? Journal of Insect Physiology 57, 1085–1089.

Heath, J. E., Hanegan, J. L., Wilkin, P. J. and Heath, M. S. (1971). Adaptation of the Thermal Responses of Insects. American Zoologist 11, 147–158.

Hellemans, S., Deligne, J., Roisin, Y. and Josens, G. (2020). Phylogeny and revision of the ‘*Cubitermes* complex’ termites (Termitidae: Cubitermitinae). Systematic entomology 46, 224–238.

Hijmans, R. J., Etten, J. van, Sumner, M., Cheng, J., Bevan, A., Bivand, R., Busetto, L., Canty, M., Forrest, D., Ghosh, A., et al. (2020). raster: Geographic Data Analysis and Modelling. https://CRAN.R-project.org/package=raster.

Huey, R. B. and Kingsolver, J. G. (1989). Evolution of thermal sensitivity of ectotherm performance. Trends in Ecology & Evolution 4, 131–135.

Huey, R. B., Kearney, M. R., Krockenberger, A., Holtum, J. A. M., Jess, M. and Williams, S. E. (2012). Predicting organismal vulnerability to climate warming: roles of behaviour, physiology and adaptation. Philosophical Transactions of the Royal Society B: Biological Sciences 367, 1665– 1679.

Johnson, M., Zaretskaya, I., Raytselis, Y., Merezhuk, Y., McGinnis, S. and Madden, T. L. (2008). NCBI BLAST: a better web interface. Nucleic Acids Research 36, W5–W9.

Jones, J. C. and Oldroyd, B. P. (2006). Nest thermoregulation in social insects. In Advances in Insect Physiology (ed. Simpson, S. J.), pp. 153–191. Academic Press.

Joseph, G. S., Seymour, C. L., Cumming, G. S., Cumming, D. H. M. and Mahlangu, Z. (2014). Termite mounds increase functional diversity of woody plants in African savannas. Ecosystems 17, 808–819.

Jouquet, P., Bottinelli, N., Shanbhag, R. R., Bourguignon, T., Traoré, S. and Abbasi, S. A. (2016). Termites: The neglected soil engineers of tropical soils. Soil Science 181, 157–165.

Kaspari, M., Clay, N. A., Lucas, J., Yanoviak, S. P. and Kay, A. (2015). Thermal adaptation generates a diversity of thermal limits in a rainforest ant community. Global Change Biology 21, 1092– 1102.

Kearney, M. and Porter, W. (2009). Mechanistic niche modelling: combining physiological and spatial data to predict species’ ranges. Ecology Letters 12, 334–350.

Kearney, M., Simpson, S. J., Raubenheimer, D. and Helmuth, B. (2010). Modelling the ecological niche from functional traits. Philosophical Transactions of the Royal Society B: Biological Sciences 365, 3469–3483.

Kellermann, V., Overgaard, J., Hoffmann, A. A., Flojgaard, C., Svenning, J.-C. and Loeschcke, V. (2012). Upper thermal limits of *Drosophila* are linked to species distributions and strongly constrained phylogenetically. Proceedings of the National Academy of Sciences 109, 16228– 16233.

Kingsolver, J. G. and Gomulkiewicz, R. (2003). Environmental variation and selection on performance curves. Integrative and Comparative Biology 43, 470–477.

Korb, J. (2003). Thermoregulation and ventilation of termite mounds. Naturwissenschaften 90, 212– 219.

Korb, J. and Linsenmair, K. E. (2000). Thermoregulation of termite mounds: What role does ambient temperature and metabolism of the colony play? Insectes sociaux 47, 357–363.

Legendre, F., Whiting, M. F., Bordereau, C., Cancello, E. M., Evans, T. A. and Grandcolas, P. (2008). The phylogeny of termites (Dictyoptera: Isoptera) based on mitochondrial and nuclear markers: Implications for the evolution of the worker and pseudergate castes, and foraging behaviors. Molecular Phylogenetics and Evolution 48, 615–627.

Mazerolle, M. J. and Linden, D. (2019). AICcmodavg: Model Selection and Multimodel Inference Based on (Q)AIC(c). https://CRAN.R-project.org/package=AICcmodavg.

Mokany, K., Raison, R. J. and Prokushkin, A. S. (2006). Critical analysis of root: shoot ratios in terrestrial biomes. Global Change Biology 12, 84–96.

Nicholls, N. and Wong, K. K. (1990). Dependence of rainfall variability on mean rainfall, latitude, and the Southern Oscillation. Journal of Climate 3, 163–170.

Noirot, C. and Darlington, J. P. E. C. (2000). Termite nests: Architecture, regulation and defence. In Termites: Evolution, Sociality, Symbioses, Ecology (ed. Abe, T., Bignell, D. E., and Higashi, M.), pp. 121–139. Dordrecht: Springer Netherlands.

Ocko, S. A., King, H., Andreen, D., Bardunias, P., Turner, J. S., Soar, R. and Mahadevan, L. (2017). Solar-powered ventilation of African termite mounds. Journal of Experimental Biology 220, 3260–3269.

Pagani, M., Freeman, K. H. and Arthur, M. A. (1999). Late Miocene atmospheric CO2 concentrations and the expansion of C4 grasses. Science 285, 876–879.

Parkash, R., Rajpurohit, S. and Ramniwas, S. (2008). Changes in body melanisation and desiccation resistance in highland vs. lowland populations of *D. melanogaster*. Journal of Insect Physiology 54, 1050–1056.

Parr, C. L., Lehmann, C. E. R., Bond, W. J., Hoffmann, W. A. and Andersen, A. N. (2014). Tropical grassy biomes: misunderstood, neglected, and under threat. Trends in Ecology & Evolution 29, 205–213.

Pearcy, R. W. and Ehleringer, J. (1984). Comparative ecophysiology of C3 and C4 plants. Plant, Cell & Environment 7, 1–13.

Poissonnier, L.-A., Arganda, S., Simpson, S. J., Dussutour, A. and Buhl, J. (2018). Nutrition in extreme food specialists: An illustration using termites. Functional Ecology 32, 2531–2541.

Porter, E. E. and Hawkins, B. A. (2001). Latitudinal gradients in colony size for social insects: Termites and ants show different patterns. The American Naturalist 157, 97–106.

Porter, W. P. and Kearney, M. (2009). Size, shape, and the thermal niche of endotherms. PNAS 106, 19666–19672.

R Core Team (2019). R: A language and environment for statistical computing. Vienna, Austria.

Rajpurohit, S., Parkash, R. and Ramniwas, S. (2008). Body melanization and its adaptive role in thermoregulation and tolerance against desiccating conditions in drosophilids. Entomological Research 38, 49–60.

Roisin, Y. (2000). Diversity and evolution of caste patterns. In Termites: Evolution, Sociality, Symbioses, Ecology (ed. Abe, T., Bignell, D. E., and Higashi, M.), pp. 95–119. Dordrecht: Springer Netherlands.

Scott Turner, J. (1994). Ventilation and thermal constancy of a colony of a southern African termite (*Odontotermes transvaalensis*: Macrotermitinae). Journal of Arid Environments 28, 231–248.

Simon, M. F. and Pennington, T. (2012). Evidence for Adaptation to Fire Regimes in the Tropical Savannas of the Brazilian Cerrado. International Journal of Plant Sciences 173, 711–723.

Singh, K., Muljadi, B. P., Raeini, A. Q., Jost, C., Vandeginste, V., Blunt, M. J., Theraulaz, G. and Degond, P. (2019). The architectural design of smart ventilation and drainage systems in termite nests. Science Advances 5, eaat8520.

Solofondranohatra, C. L., Vorontsova, M. S., Hackel, J., Besnard, G., Cable, S., Williams, J., Jeannoda, V. and Lehmann, C. E. R. (2018). Grass functional traits differentiate forest and savanna in the Madagascar Central Highlands. Frontiers in Ecology & Evolution 6, 184.

Sternberg, L. D. S. L. (2001). Savanna–forest hysteresis in the tropics. Global Ecology and Biogeography 10, 369–378.

Still, C. J., Pau, S. and Edwards, E. J. (2014). Land surface skin temperature captures thermal environments of C3 and C4 grasses. Global Ecology and Biogeography 23, 286–296.

Støen, O.-G., Okullo, P., Eid, T. and Moe, S. R. (2013). Termites facilitate and ungulates limit savanna tree regeneration. Oecologia 172, 1085–1093.

Sunday, J. M., Bates, A. E. and Dulvy, N. K. (2011). Global analysis of thermal tolerance and latitude in ectotherms. Proceedings of the Royal Society B: Biological Sciences 278, 1823–1830.

Sunday, J. M., Bates, A. E., Kearney, M. R., Colwell, R. K., Dulvy, N. K., Longino, J. T. and Huey, R. B. (2014). Thermal-safety margins and the necessity of thermoregulatory behavior across latitude and elevation. PNAS 201316145.

Tang, H., Armston, J., Hancock, S., Marselis, S., Goetz, S. and Dubayah, R. (2019). Characterizing global forest canopy cover distribution using spaceborne lidar. Remote Sensing of Environment 231, 111262.

Traniello, J. F. A. and Leuthold, R. H. (2000). Behavior and ecology of foraging in termites. In Termites: Evolution, Sociality, Symbioses, Ecology (ed. Abe, T., Bignell, D. E., and Higashi, M.), pp. 141–168. Dordrecht: Springer Netherlands.

Turner, J. S. (2001). On the mound of Macrotermes michaelseni as an organ of respiratory gas exchange. Physiological and Biochemical Zoology 74, 798–822.

Turner, J. S. (2019). Termites as mediators of the water economy of arid savanna ecosystems. In Dryland Ecohydrology (ed. D’Odorico, P., Porporato, A., and Wilkinson Runyan, C.), pp. 401–414. Cham: Springer International Publishing.

Vesala, R., Harjuntausta, A., Hakkarainen, A., Rönnholm, P., Pellikka, P. and Rikkinen, J. (2019). Termite mound architecture regulates nest temperature and correlates with species identities of symbiotic fungi. PeerJ 6, e6237.

Wiafe, E. D. (2016). Tree Species Composition of Kakum Conservation Area in Ghana. Ecology and Evolutionary Biology 1, 14.

Willmer, P. G. (1982). Microclimate and the environmental physiology of insects. In Advances in Insect Physiology (ed. Berridge, M. J., Treherne, J. E., and Wigglesworth, V. B.), pp. 1–57. Academic Press.

Wood, T. G. and Thomas, R. J. (1989). The mutualistic association between Macrotermitinae and Termitomyces. In Insect-Fungus Interactions, pp. 69–80 p. Academic Press.

Woon, J. S., Boyle, M. J. W., Ewers, R. M., Chung, A. and Eggleton, P. (2018). Termite environmental tolerances are more linked to desiccation than temperature in modified tropical forests. Insectes sociaux 66, 57–64.

